# Uniformity of performance during the collection of maximum voluntary contraction tasks for the muscles of the forearm

**DOI:** 10.1101/2023.03.29.534788

**Authors:** Mercedes Aramayo Gomes Rezende, Oluwalogbon O Akinnola, Angela E Kedgley

**Author notes:** Corresponding author: Angela Kedgley.

## Abstract

Electromyographic (EMG) signals are used to gain insight into muscle activation patterns and thus neuromuscular control. To allow for comparisons between studies and participants, the EMG signal is generally normalised, with the signal obtained during a task that is designed to elicit maximum voluntary contraction (MVC) frequently used as the basis. Recommendations for how to collect MVCs have been made; however, previous studies have not been able to determine if the EMG variation noted within a population was due to different muscle activation patterns, or the tasks being performed differently, or other variables such as skin impedance, or the stochastic nature of electromyography.

EMG signals were recorded during hand-wrist tasks selected to elicit MVCs in the muscles of the forearm – pull up, push down, radial pull, ulnar pull, pull, pronation, finger flexion, finger extension, and grip – as well as two activities of daily living – pouring a glass of water from a jug and turning a key in a lock. A load cell mounted to a statically mounted handle was used to record contemporaneously the forces and moments exerted by participants in pull up, push down, radial pull, ulnar pull, pull, pronation, finger flexion, and finger extension tasks.

Ninety percent of tasks yielded the expected load cell outputs for the directed tasks and thus were considered as having been performed correctly. The tasks performed incorrectly were not the same for all participants, nor were they all performed by the same participants. Of note was that there were instances when a task was performed incorrectly but still an expected MVC was achieved. The EMG signals showed similar variation to that seen in previous studies. However, the applied forces and moments did not appear to explain the variation seen in the tasks that elicited MVCs.

The results of this study indicate that different muscle activation patterns may be used to exert the same force by the hand. Thus, it may not be possible for a given task to elicit MVC in the same muscle in all people. However, by using several activities, MVCs for the forearm muscles may be obtained for most of the population. Beyond designing EMG protocols, the results of this study suggest that people have unique muscle activation patterns and raise questions as to whether this is a result of physiology or conditioning.

## 1. Introduction

Electromyography (EMG) is commonly used to observe muscle activity and has many different applications in the medical field, including as an assessment tool to evaluate muscle and motor neuron health (Reaz et al., 2006). However, EMG is sensitive to changes in conditions such as electrode placement and sweat on the skin (Abdoli-Eramaki et al., 2012). To enable inter- or intra-subject comparisons, measurements of muscle activity during maximum voluntary contraction (MVC) are one means used to normalize EMG data (Besomi et al., 2020). Using signals during MVC to normalise EMG data also provides insight into the effort exerted by a participant. Historically, no standard protocol for the collection of MVCs for the muscles of the forearm existed. Thus, inter-study comparisons were limited, as the data were normalised by values that were potentially very different, resulting in various inferences on relative effort. This is often a risk when comparing EMG results, but a standard protocol for the collection of MVC can provide mitigation. Recommendations on best practice to collect MVC of the forearm muscles have been made; Ngo and Wells (2016) found that using a series of resisted moment exertions would elicit greater muscle activity than the commonly used power grip task. Akinnola et al. (2020) presented tasks most likely to elicit MVC in muscles of the forearm and suggested that targeted protocols could be produced for efficient collection of MVCs. However, a limitation of the study was that it was not possible to verify that participants performed the tasks as instructed. Thus, the observed variation in the tasks that elicited MVC in each muscle could have been the result of participants performing different actions, rather than physiological differences in anatomy or activation patterns.

To create an efficient protocol, it is important to understand the relationship between the forces and moments applied by participants and their muscle activity during tasks designed to elicit MVCs. Thus, it can be determined whether each task is performed as intended by participants, increasing confidence that the recommended tasks will result in the MVC of specific muscles. The aims of this study were to build on the work of Akinnola et al. (2020) by quantifying EMG signals and kinetics during a series of isometric tasks to verify the recommendations for a standardised protocol for the collection of MVCs in muscles of the forearm and to understand the source of any inter-participant variability.

## 2. Materials and methods

Twenty-two right-handed participants (14 female, 8 male, 24.5 ± 4.96 years old) took part in the study. Ethical approval was acquired from the Imperial College Research Ethics Committee. All participants provided written informed consent.

Nine surface EMG sensors (Delsys Trigno, Natick, MA, USA) were placed on the participant’s dominant forearm to capture muscle activity of the flexor carpi radialis (FCR), flexor digitorum superficialis (FDS), flexor carpi ulnaris (FCU), extensor digitorum communis (EDC), extensor carpi ulnaris (ECU), extensor carpi radialis (ECR), pronator teres (PT), biceps and triceps. The signals from the biceps and triceps were only employed to ensure that participants performed the activities using their hands and forearms, rather than relying on these larger muscles. One sensor was secured over the belly of each muscle using double-sided tape. It was ensured that electrodes did not contact one another during forearm movement to avoid unnecessary noise generation. Prior to starting the task, crosstalk between electrodes was examined to minimise it. Participants were then seated on an adjustable stool facing a table where a handle was mounted (Figure 1). The handle was attached to a six degree of freedom load cell (M3944, Sunrise Instruments, Canton, MI, USA) that was used to measure the forces and moments applied by the participant during each task.

**Figure 1:**
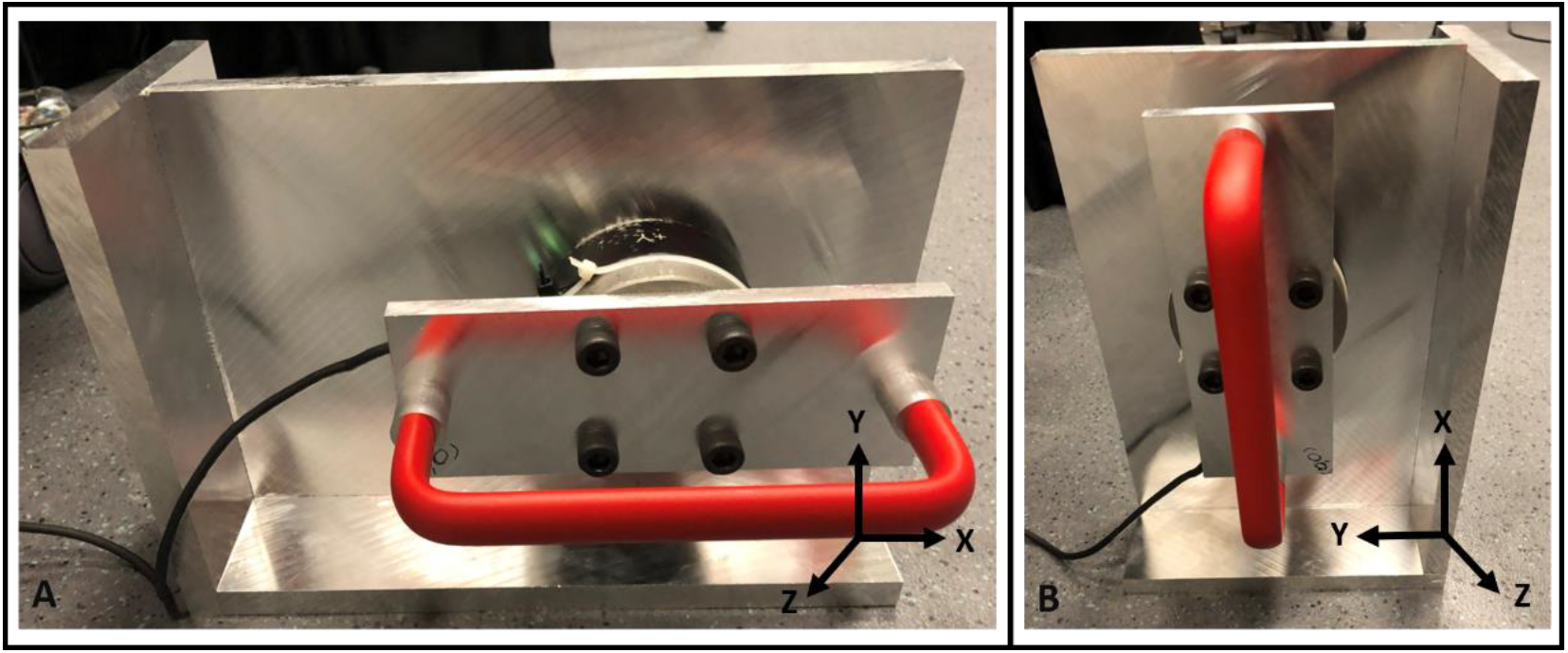
Handle position and associated axes. (A) For the pull up and push down tasks the rig handle was oriented horizontally. (B) For the radial pull, pull, ulnar pull, pronation, finger flexion and finger extension tasks the rig handle was oriented vertically. This enabled the tasks to be performed in accordance with the protocol developed by Akinnola et al. (2020). To change the orientation of the handle, the handle, load cell, and associated base were rotated in their entirety and clamped to the table.

Each participant was then asked to perform a series of tasks, adapted from Ngo and Wells (2016), that were designed to elicit MVC in each muscle. These tasks were pull up, push down, radial pull, ulnar pull, pull, pronation, finger flexion, finger extension, grip (Akinnola et al., 2020), as well as two activities of daily living (ADL), which were pouring a glass of water from a jug and turning a key in a lock. The ADL tasks provided a baseline measure for expected muscle activations. Participants sat on a height-adjustable stool with the rig mounted on a table in front of them. For the tasks involving the rig, participants were asked to keep their elbows on the table throughout the task. Each task was explained to the participants, and they were encouraged to try each before recording. Tasks were performed once for a period of 5 seconds. Verbal encouragement was given throughout the 5 seconds so that the participants felt compelled to do their very best. Participants were allowed to rest for a minimum of two minutes between tasks.

For all tasks, participants sat with their feet flat on the floor. For all but the pull up and push down tasks the rig was orientated vertically; participants gripped the bar with the wrist in neutral position and tried to rotate the beam by 90° in the appropriate anatomical direction. Participants did not grip the bar during the finger tasks, instead they pressed their extended fingers, from the proximal interphalangeal joint distally, against the bar and then pushed in the appropriate direction. For the grip task, participants sat with their elbow flexed 90°, and their shoulder in a neutral position. They squeezed a dynamometer as hard as they could by only flexing their fingers. For the ADL tasks, participants started and finished each task with their dominant hand flat on the table; they could take as long has needed to complete the tasks. For the ADL pouring task, they picked up a 1.5 L jug filled with water, poured approximately 250 mL of water into a cup, stabilising the cup using their non-dominant hand if needed, and returned the jar to its original location. For the ADL key task, participants turned a key that was already placed in a lock anticlockwise until its maximum position and then released it.

Custom code written in MATLAB (MathWorks, Natick, MA, USA) was used to analyse the data from the EMG sensors and the load cell. The EMG signal had the DC offset removed, was rectified, and then filtered using a 2^nd^ order low pass filter with a cut-off frequency of 13 Hz (Robertson and Dowling, 2003). The activity of each muscle in each task was noted and the activity during MVC for a given muscle was that which elicited the peak across all tasks. The task was identified and all remaining EMG data for that muscle were normalised by the MVC.

Load cell data were passed through a zero-phase filter. The peak force and moment in each degree of freedom (DoF) across all tasks was used to normalise the data. The force and moment patterns were compared visually to establish if the task had been correctly performed and to determine whether there were any repeating patterns, such whether high values in a primary DoF were accompanied by peaks secondary DoFs. The primary DoF was that in which the greatest force or moment for a given task was expected; for example, the positive y-direction for the pull up task (Figure 1A) or positive z-direction in the pull task (Figure 1B). Forces and moments in DoFs other than the primary were designated as secondary. Tasks were determined as being performed correctly if the direction and magnitude of the force or moment in the primary DoF of the given task were as expected.

Non-parametric Friedman tests were used to test for statistical differences between the muscle activities during the tasks; significance was defined as p < 0.05.

## 3. Results

The distribution of tasks that produced MVC varied considerably for each muscle, with no muscle always generating MVC in a single task and no task producing MVC for every muscle (Figure 2A). For some muscles, only one participant produced MVC in a specific task. Replacing those tasks with the tasks that elicited the second highest activity for that muscle reduced the number of tasks used to generate MVC for each muscle (Figure 2B).

**Figure 2:**
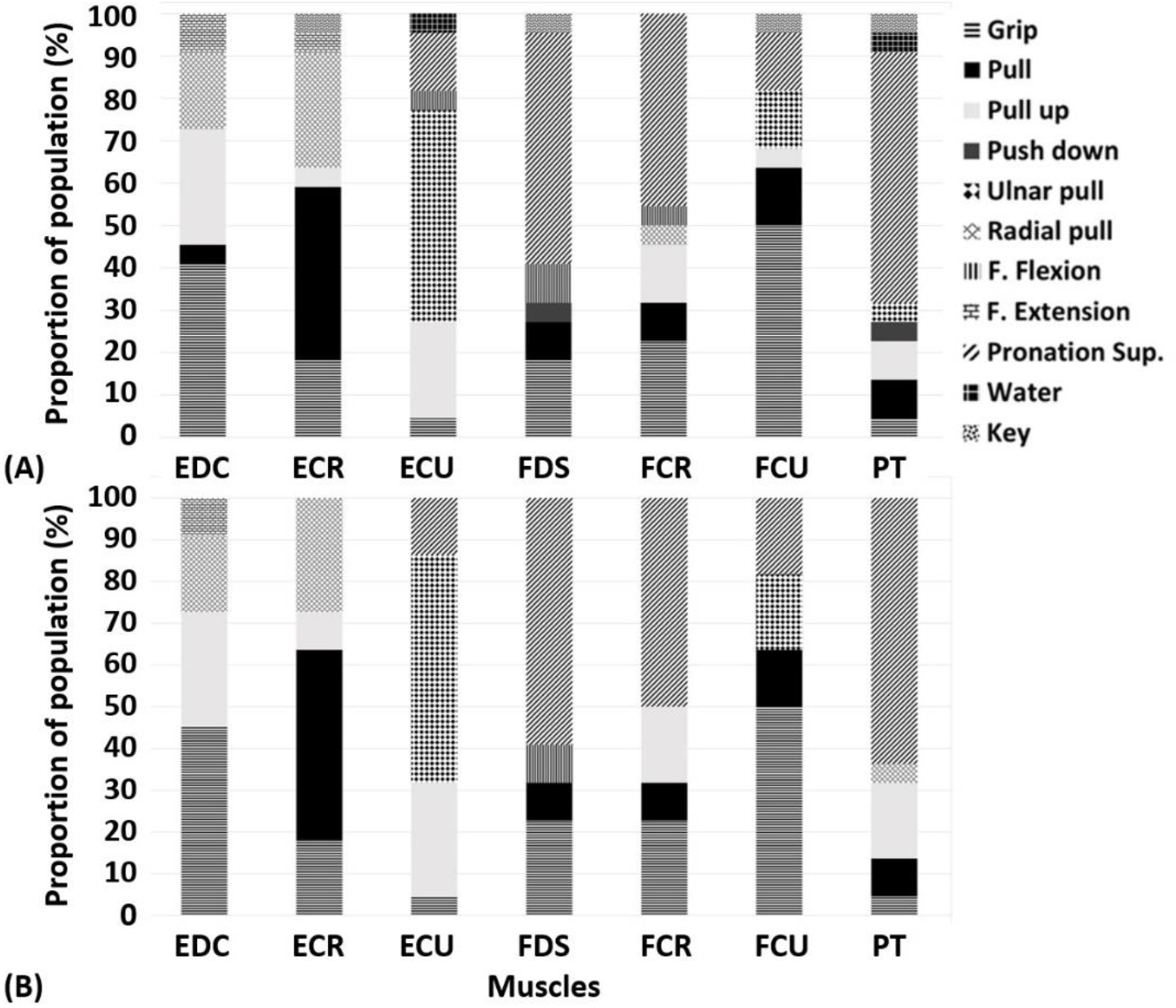
Distribution of the participants that generated maximum voluntary contraction. (A) considering all tasks and (B) upon replacement of those tasks which generated MVC for a given muscle in only one participant with the task that produced the next greatest level of activity for that participant for the extensor digitorum communis (EDC), extensor carpi radialis (ECR), extensor carpi ulnaris (ECU), flexor digitorum superficialis (FDS), flexor carpi radialis (FCR, flexor carpi ulnaris (FCU), and pronator teres (PT) muscles.

For some muscles there was no activity that clearly elicited more activity than that produced by other tasks. An example of this is the FCR (Figure 3A). In contrast, for other muscles differences were noted in the muscle activities within the tasks (p < 0.001); for example, pronation was most likely to produce MVC for the PT (Figure 3C; p < 0.02).

**Figure 3:**
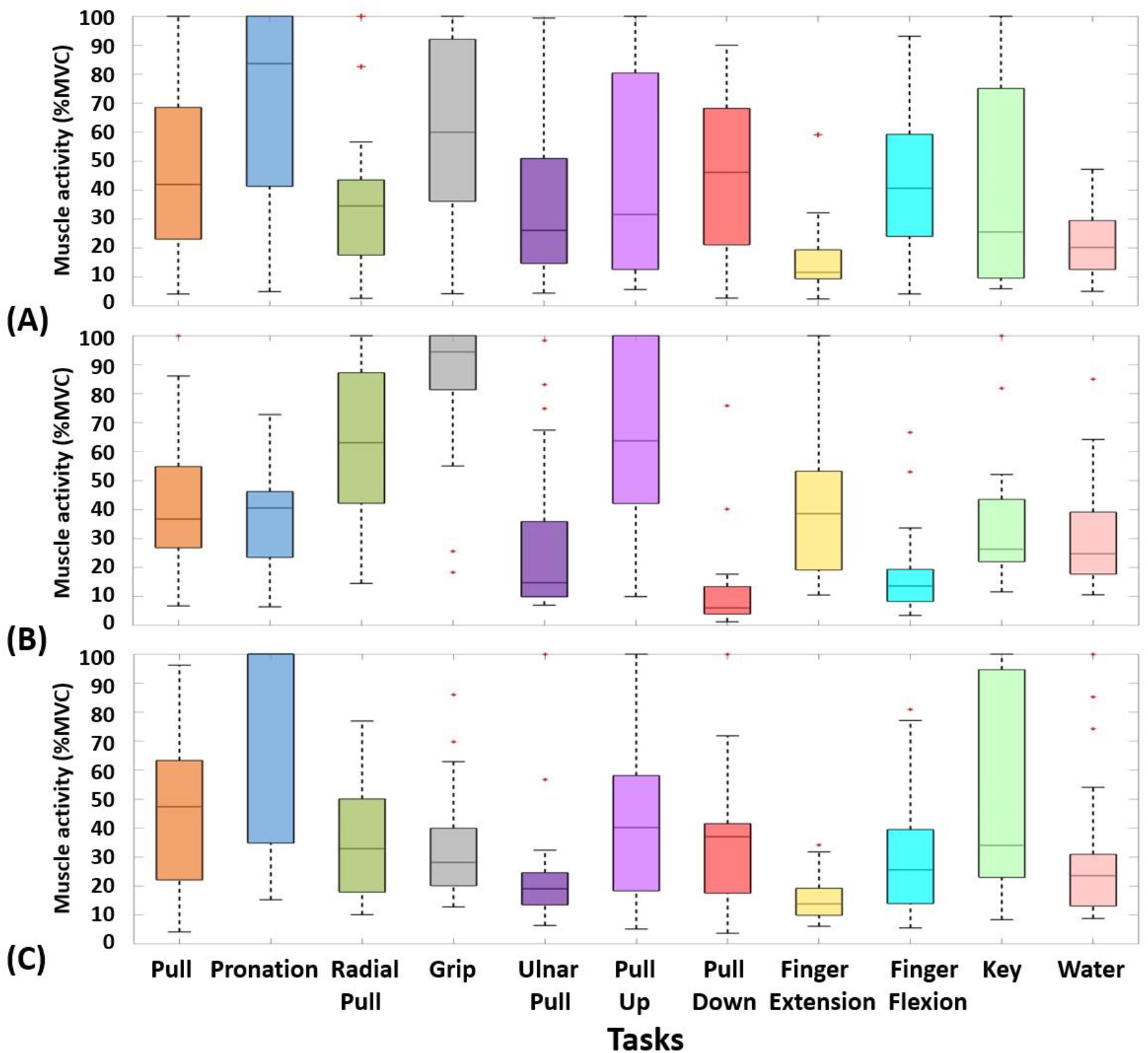
Muscle activity in all nine tasks, expressed as a percent of maximum voluntary contraction. (A) Flexor carpi radialis, (B) extensor digitorum communis, and (C) pronator teres.

The force data were used to verify that the tasks were performed correctly. The majority of participants performed each task correctly by generating force and moments in the required primary DoF (Figure 4). For example, the largest moment about the z-axis of the handle was generated during the pronation task (p < 0.004) and the forces and moments in the primary DoF were significantly greater higher than in the other DoFs (p < 0.001).

**Figure 4:**
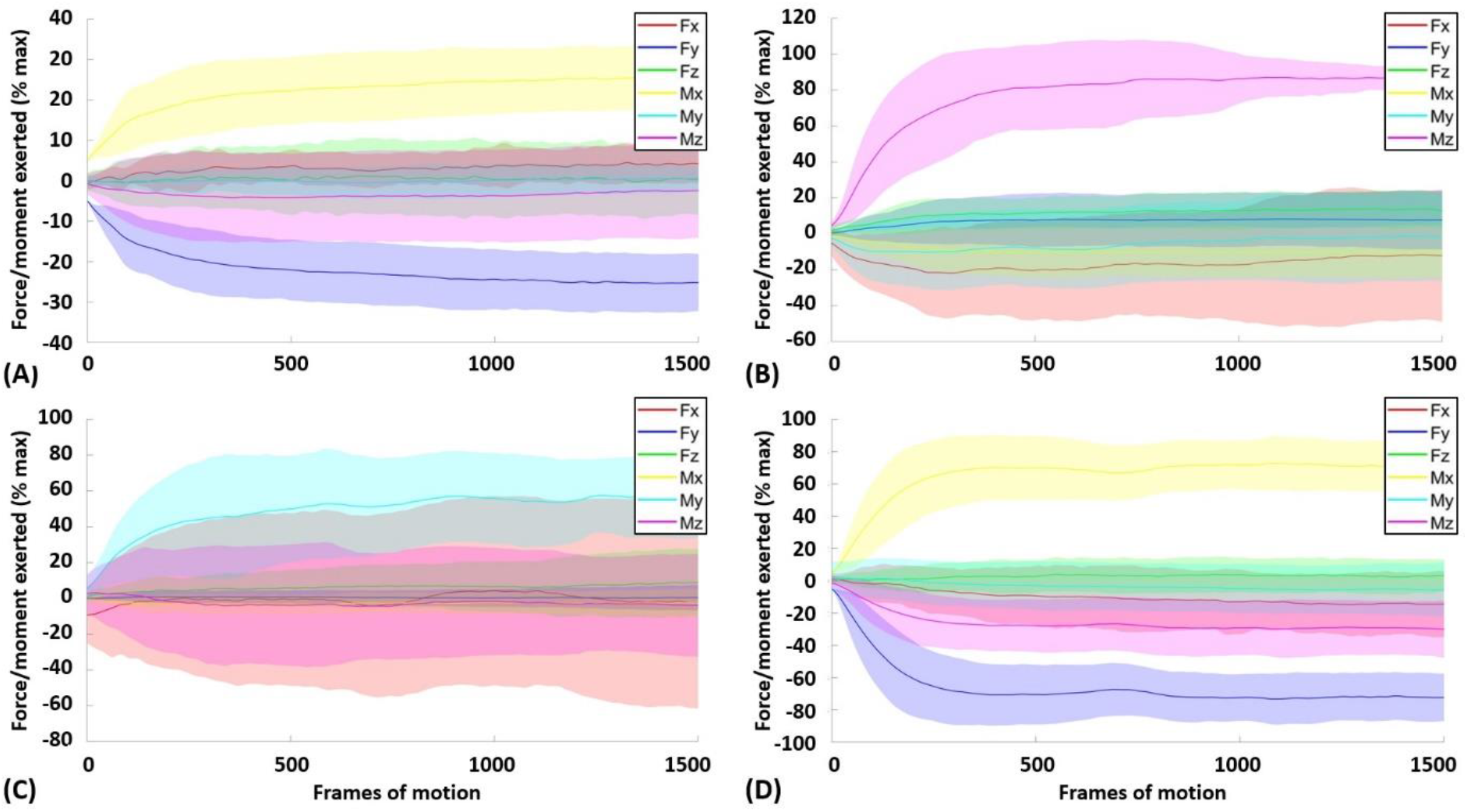
Mean ± one standard deviation of the forces and moments exerted by participants during different tasks. (A) Finger extension: with the handle in the vertical position, participants were asked to extend their fingers against it. Forces were expected to be negative in the y-axis with potential for a positive moment about the x-axis. (B) Pronation: with the handle in the vertical position, participants were asked to try to rotate the handle anti-clockwise. Moments were expected to be positive about the z-axis. (C) Ulnar pull: with the handle in the vertical position, participants were asked to rotate their wrist towards their little finger. Moments were expected to be positive about the y-axis. (D) Pull down: with the handle in the horizontal position, participants were asked to pull the handle straight down. Forces were expected to be negative in the y-axis with potential for a positive moment about the x-axis.

Only 10% of tasks were performed incorrectly. No task was performed correctly by every participant and the participant(s) who performed a task incorrectly varied for the different tasks. Qualitative differences in secondary moment and force patterns for each participant were observed, but the primary moments and forces for each specific task were always exerted in the same direction with differences only in the amplitude.

## 4. Discussion

No one task elicited MVC for a given muscle in every person nor did any task produce MVC in every muscle. Some tasks resulted in only one participant producing MVC for a given muscle. When these were replaced with the task that produced the second highest activity, more uniformity in the tasks was observed. This suggests that while tasks may target specific groups of muscles, different people have different combinations of muscles they use to perform a given task. Previous research has shown that muscles are activated when a task aligns with their anatomical function (Greig and Wells, 2008) and these data may corroborate this.

The load cell data showed that, overall, tasks were performed correctly by participants. The uniformity seen in the forces in the primary DoF of the tasks suggests that participants generally understood the directions given and could follow them. Thus, it can be hypothesised that the variety in the MVCs elicited by a task is not necessarily the result of the task being performed incorrectly. Even when participants were identified as being outliers when considering the activity for a given muscle in a task, they were found to be performing the task correctly. In fact, there were only two instances where participants identified as outliers in a task performed the task incorrectly. This corroborates the EMG data in suggesting that different combinations of muscle activity can be used to achieve a task. Previous literature has found similar results: Abernethy et al. (1990) found high inter-subject variability in upper limb muscle activation during a golf swing and Takamori et al. (2020) found variations in forearm muscle use during pronation. Participants were not consistently identified as outliers across all tasks, nor did they achieve MVC in the same muscles. This indicates that there was not an alternate muscle activation by which to group them. Of note, however, is that a participant can perform a task incorrectly yet still elicit MVC in the intended muscle. For example, one participant performed the radial and ulnar pull tasks incorrectly. However, they still generated MVC values during those tasks in the FCR and ECU, respectively. Conversely, another participant did not generate much force during the flexion task, yet demonstrated near MVC activity in the FDS. This demonstrates that the muscles can be very active without generating much force on the surrounding environment. Co-contraction, for example, could result in very high muscle activity, as the agonists work against the antagonists, but with little force being exerted on the handle. These may seem to suggest that prescribed tasks are unreliable for eliciting MVC, but cases like those mentioned above were in the minority and, furthermore, were unpredictable. Therefore, there is still benefit in having a standardised protocol that captures MVCs for the majority of the population.

Variation was seen in the secondary forces and moments, and it was thought that this could explain the variation in the muscles that produced MVC, as the relative magnitudes were similar. For example, if a participant also extended their wrist during radial pull, we would expect MVC for the ECR but perhaps not for the FCR. Thus, one would expect that participants who produced MVC in the same task would have similar secondary forces and moments. However, this was not seen consistently among participants. For example, participants A and B produced the MVC for the ECU in the ulnar pull task, participant C produced MVC in the pull up task, and participant D produced MVC in the flexion task. However, the force patterns in the six DoFs for all four participants during the ulnar pull task were similar and, when differences were noted, there was more similarity between the force patterns of participants B and C than between A and B. This points to the fact that the generated force does not perfectly relate to observed muscle activity. Participants A, B, and C all produced the second highest activity for the ECU during the extension task. Participant D produced their second highest activity during grip. Yet, there was little difference between participant D and the participants A-C during the extension task. The data indicate that it is not differences in the force being generated that are the cause of the differences in the observed EMG.

As participants performed the tasks correctly but variation was still seen in the MVCs, which was not explained by variation in the forces, it is hypothesised that muscle activation patterns vary between people. The data show that muscles anatomically able to contribute to a task are generally maximally activated during that task but variations may be because of physiological differences between participants. These differences could be the result of participant-specific anatomy or lifestyle differences, such as sporting activities or playing a musical instrument.

As is common with investigations relating to MVC, one limitation of this study was that it is difficult to ascertain whether MVC truly was generated. This would impact the results, as a task that may be better at producing MVC in a given muscle may not appear so, if participants did not exert maximum effort. To ameliorate this: participants were given time to practice, ensuring they understood the motion; verbal encouragement was given; and participants rested between tasks to limit fatigue. Further, it could be argued that a participant’s ability to interpret a task and perform it correctly is part of its viability on being used in a protocol. If a task was anatomically advantaged in producing MVC but participants repeatedly do not achieve the MVC, it would indicate that some change is needed. Fortunately, this was not the case in this study.

Another limitation was that many of the tasks involved gripping as well as force generation in a given direction. As muscles like the FDS and EDC are bi-articular, other muscles involved in a motion may not have needed to be as active and thus underperformed in the task anatomically suited to them. It has been shown that resisted moment tasks produce higher activation but perhaps a different mechanism of resisting the hand, perhaps the use of elastic, could better target muscles, allowing for more precise protocols.

## 5. Conclusion

The data collected in this study indicate that a protocol can be designed to obtain MVC for the forearm muscles for the majority of the population. The force data confirms that participants performed the tasks correctly and indicates that the variations in the secondary forces and moments were not the reason for the differences in the tasks that elicited MVC. It is hypothesised that physiological differences could explain why different muscles are maximally activated doing the same task. This has implications for understanding and modelling activation patterns for the muscles of the forearm.

